# Sephin1 alleviates white matter injury by protecting oligodendrocyte after intracerebral hemorrhage

**DOI:** 10.1101/2025.06.26.661862

**Authors:** Fangyu Liu, Qianqian Ma, Xinhui Li, Zuoqiao Li, Peng Wang, Wensong Yang, Yiqing Shen, Peizheng Li, Mingjun Pu, Zhongsong Xiao, Peng Xie, Qi Li

## Abstract

**Background:** White matter injury (WMI) caused by intracerebral hemorrhage (ICH) is a major neuropathological feature closely associated with neurological impairments such as motor and sensory dysfunction. Oligodendrocytes (OLs), which are responsible for repairing WMI, also suffer severe death resulting from the compression of hematoma and secondary neuroinflammation after ICH. Sephin1, a selective inhibitor of PPP1R15A, has been shown to reduce general protein synthesis and protect OLs by prolonging the integrated stress response (ISR). We aimed to evaluate the effectiveness of Sephin1 in protecting OLs in experimental ICH mice and primary OLs and microglia co-cultures.

**Methods:** We first determined the performance of ICH mice treated with Sephin1 or vehicle in multiple behavioral tests. To investigate dynamic changes in the number of OLs surrounding the hematoma after ICH, we labeled and tracked apoptotic, proliferating, and mature OLs using immunofluorescence staining.

**Results:** Sephin1 treatment improved long-term neurological function after ICH, which was accompanied by a significant alleviation of WMI in the perihematomal region. Our data indicated that Sephin1 dramatically increased the population of OLs in the perihematomal region after ICH by inhibiting OL apoptosis and promoting OL proliferation. Moreover, Sephin1 treatment attenuated neuroinflammation after ICH by inhibiting microglial polarization to the M1 phenotype. *In vitro*, a co-culture model of primary OLs and microglia demonstrated that Sephin1 preserved the viability of OLs under pro-inflammatory conditions.

**Conclusions:** Our observations suggest that Sephin1 is a promising therapeutic drug to preserve the OLs and alleviate WMI around the hematoma in ICH, highlighting its translational potential to improve long-term neurological recovery in hemorrhagic stroke.

## Introduction

Intracerebral hemorrhage (ICH) is the most serious form of stroke, accounting for approximately 15% of all subtypes, and has high mortality and disability rates.^1,2^ Despite recent advances in ICH treatment, there are few interventions to effectively improve the prognosis and clinical outcomes of ICH.^3,4^ Therefore, novel treatment strategies that address the high disability rate of ICH and improve long-term neurological function are urgently required.

Proximal or distal white matter injury (WMI) caused by ICH is closely associated with poor prognosis, especially motor and sensory dysfunction.^5^ More than 77% of patients with ICH experience WMI.^6^ However, effective therapies for WMI following ICH require further investigation.

Oligodendrocytes (OLs), which are responsible for myelination in the central nervous system (CNS), play an important role in the remyelination and neurological functional repairments of ischemic and hemorrhagic stroke.^7,8^ Primary injury from direct compression and barotrauma caused by hematoma, and secondary injury from neuroinflammation and hematotoxic products after ICH, lead to severe OLs death following apoptosis, ferroptosis, and necrosis.^9–11^ The contributing factor to this phenomenon is that OLs (including oligodendrocyte progenitor cells [OPCs] and mature OLs) are vulnerable to cytotoxic and excitotoxic factors because of their high metabolic rates, lipid richness, and lack of essential glutathione and glutathione peroxidase.^12,13^ However, the density of OLs, increases markedly a few days after ICH due to proliferation, migration, or both.^14^ However, the differentiation of OPCs into myelin-producing OLs and the remyelination of injured axons by mature Ols are implicated in microglia-induced inflammatory responses.^15^ Therefore, protecting OLs from primary and secondary injuries induced by ICH may be beneficial for the restoration of WMI and the recovery of neurological function.

The integrated stress response (ISR) is a cytoprotective mechanism activated by eukaryotic cells to restore cellular homeostasis in response to various intrinsic and extrinsic stressors.^16,17^ ISR can be triggered in many CNS pathologies such as ischemic stroke, multiple sclerosis, and hypoxia-induced diffuse WMI.^18–20^ Similarly, tissues in the perihematomal region undergo various pathological changes after ICH, including endoplasmic reticulum stress, oxidative stress, neuroinflammation, and nutritional deprivation, all of which are potent inducers of ISR.^21^ The central event of ISR system is the phosphorylation of eukaryotic translation initiation factor 2 (eIF2α), which inhibits the activity of eukaryotic translation initiation factor 2B (eIF2B) and results in rapid attenuation of global protein synthesis, while selectively favoring the translation of selected genes implicated in the stress response.^22^ Subsequently, the activation of ISR could conserve cellular resources, restore cellular homeos Ki67 tasis, and promote cell survival.^23^ Prolonging the ISR can protect remyelinating OLs and promote remyelination in an inflammatory CNS environment.^24^ Additionally, the regulating ISR could exert anti-inflammatory effects on microglia via promoting M2 polarization.^25^

Sephin1, a selective inhibitor of PPP1R15A, specifically inhibits GADD34-PP1c-mediated dephosphorylation of p-eIF2a, thereby prolonging the ISR, which has been used to treat a variety of animal models of neurological diseases.^26–29^ However, the potential effects of Sephine1 on ICH remain unknown.

We, therefore, aimed to establish ICH models in mice by injecting bacterial collagenase type VII-S into the right basal ganglia and to explore the effects and mechanisms of Sephin1 post-ICH. We found that Sephin1 significantly protected OLs and promoted the repair of the WMI surrounding the hematoma after ICH by prolonging the ISR. Furthermore, in an *in vitro* experiment, apoptosis of OLs co-cultured with hemoglobin-activated microglia was attenuated by Sephin1. Together, this evidence demonstrates that Sephin1 could improve the potential repair of WMI and promote long-term neurological recovery after ICH by preserving the OLs around the hematoma.

## Methods

### Data availability

Data supporting the findings of this study are available from the corresponding author upon reasonable request. Supplementary material is available at Stroke online. The materials are available from commercial sources.

### Animals

All animal experiments were designed in accordance with the ARRIVE guidelines. All experimental procedures were approved by the Ethics Committee of Chongqing Medical University. Generally, eighteen-month-old male C57BL/6J mice (27–33 g) were obtained from Chongqing Medical University Laboratory Animal Centre (Chongqing, China). Mice were housed under specific pathogen-free conditions with a 12/12-h light/dark cycle, controlled temperature and humidity, and *ad libitum* access to food and water.

The sample size for each analysis in this study was determined by power calculation (α=0.05, power=0.8). In addition, the mean difference and standard deviation were based on the pre-experiment we performed. Then, we calculated that a sample size of n=17 would be needed to capture the treatment effects at each time point. In consideration of mortality in experimental animals, we added n=1 to each time point in the different groups.

A total of 216 mice were used in this study and randomly assigned to three groups: Sham, ICH + Vehicle, or ICH + Sephin1 groups (n=72 per group) using computer-generated randomization before surgical procedure. An independent researcher performed the assignment to maintain blinding. The final randomization list documented group assignments for all mice and was archived for reproducibility. Mice were excluded from the analysis if they died from surgery or didn’t present significant neurological deficits at 24 hours after surgery. Three mice (one per group) were excluded from final analyses due to perioperative mortality (death within 24 hours post-surgery). All other enrolled mice completed the full protocol and were included in analyses. The time-course of behavioral testing, pathology analysis was performed from 3 to 28 days after ICH-induced (Figure 1A).

**Figure 1.**
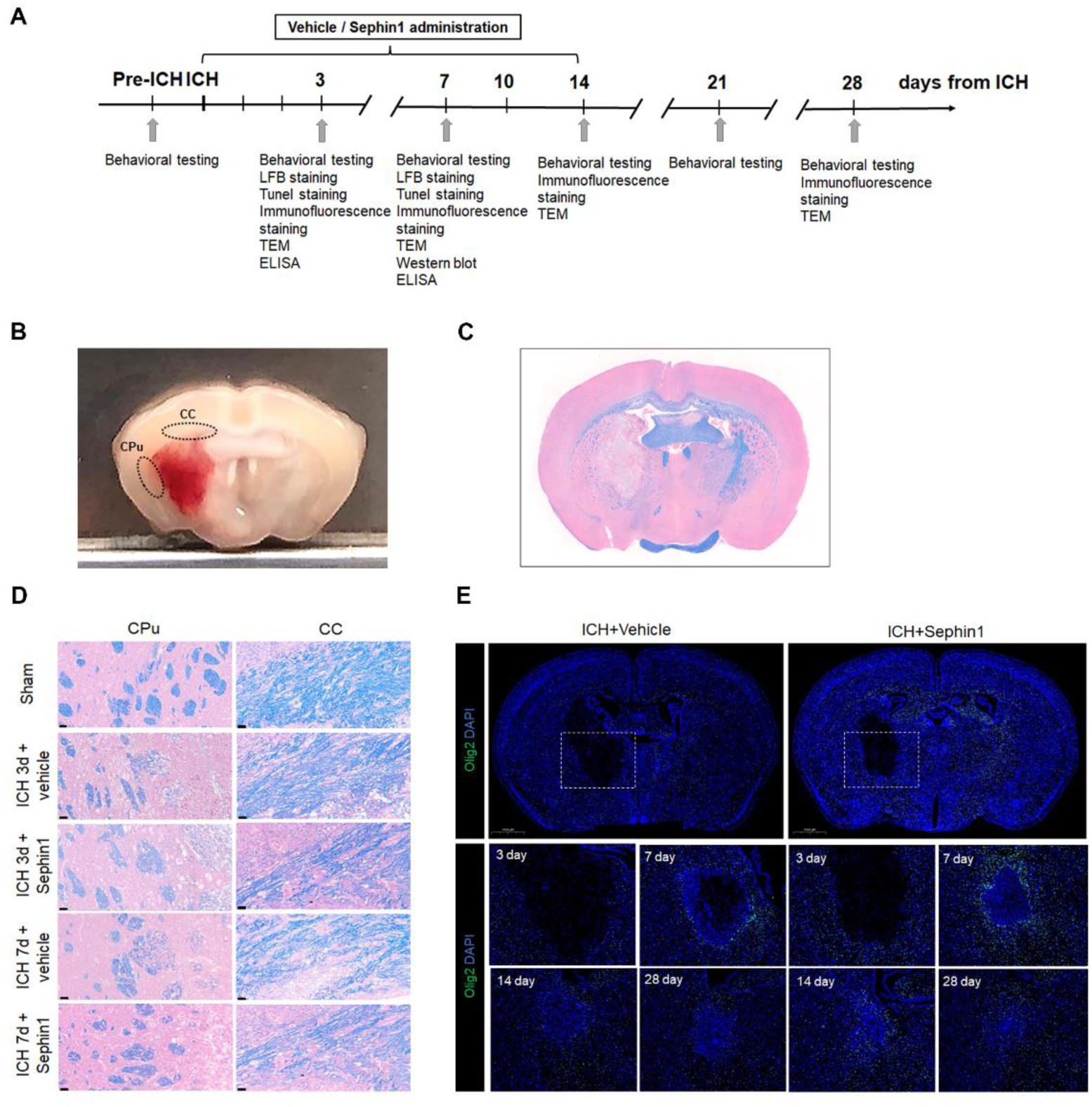
ICH leads to severe WMI and apparently decreasing the population of OLs in perihematomal region. **A,** Time schedule for the vivo portion of the experimental design. **B,** The scheme of coronal brain section illustrated the location for WMI and OLs measurements. **C,** Coronal section of brain as shown in LFB staining. **D,** LFB staining showed severe WMI in the CPu and CC surrounding the hematoma after experimental ICH in mice. Lens: × 400; Scale bar: 20 μm. **E,** Immunofluorescence staining for Olig2 and DAPI indicated the time-dependent changes in the density of OLs. Lens: × 15; Scale bar: 1000 μm.

### ICH model

To better understand WMI in the perihematomal region after ICH, we established a mouse model of ICH in which the hematoma was located below the corpus callosum (CC) and the external capsule (EC). C57BL/6 mice were anesthetized by 5% isoflurane inhalation and placed in a stereotaxic frame (Rivard, Shenzhen, China). Subsequently, the mice were maintained under anesthesia with 1.5–2% isoflurane. The core body temperature of the mice was maintained at 37.0°C with an adjustable heating pad. A 1-mm burr hole was made in the skull (0.5 mm anterior, 2 mm right lateral, relative to the bregma), and a Hamilton needle (Hamilton, Bonaduz, Switzerland) was inserted at a depth of 3.85 mm below the skull. The ICH model was induced within 5 min by injecting 0.0375 U of bacterial collagenase type VII-S (C2399, Sigma-Aldrich, St. Louis, MO) in 0.375 μL saline into the right basal ganglia using a microperfusion pump system (Rivard, Shenzhen, China). To avoid backflow, the needle was left for an additional 10 min after the injection was completed and then removed gently at a rate of 1 mm/min. Finally, the burr hole was sealed with bone wax and the incision was sterilized and sutured. In sham group, the mice underwent the same procedure, but were injected with the same volume of saline without collagenase.

### Administration of Sephin1

Sephin1 (HY-111022) was purchased from MedChemExpress (Monmouth Junction, NJ). The treatment protocol and dosage of Sephin1were based on our pre-test results (Supplementary Figure 1A-C). We performed experiments to test the effect and safety of different dose of Sephin1 (5/10/20 mg/kg *in vivo* and 20/50/100 μM *in vitro*). For the animal experiment, Sephin1 was dissolved in 1% dimethyl sulfoxide (DMSO) and double-distilled water to a final concentration of 1 mg/ml; 10 mg/kg Sephin1 or an equivalent volume of vehicle (1% DMSO in 0.9% NaCl) was administered by intraperitoneal injection daily for 3/7/14 consecutive days starting from 30 min after ICH. For *the vitro* experiments, the concentration of Sephin1 in culture medium of OLs was prepared as 50 μM.

### Behavioral testing

We performed the mNSS, accelerated rotarod test, forelimb grip assessment, hot-plate test, and balance beam walking test to evaluate neurological function before surgery and at 3/7/14/21/28 days after ICH. The tests were performed by a trained investigator who was blinded to the experimental groups. All animals received three days of behavioral training before ICH induction.

#### mNSS assessment

To assess neurological deficits after ICH, mice were scored using the mNSS, which is a complex behavioral test that includes motor, sensory, reflex, and balance tests. Neurological function was rated on a scale of 0–18 (0=normal function, 18=maximal deficit).

#### Accelerated rotarod test

For the accelerated rotarod test, mice were placed on a Six-lane Accelerating Rotarod Apparatus (SA102, Sansbio Co. Ltd., China) to assess sensorimotor coordination and fatigue resistance. The rotating rod was accelerated consistently from 4 to 40 rpm in 5 min, increasing by 8 rpm every 60 s, until the maximum speed at 300s. The test ended when the mice fell off the rods or grabbed the apparatus and spun around for two consecutive revolutions without attempting to walk on the rods. The duration (s) for which the mice stayed on the Accelerating Rotarod Apparatus was recorded. According to the experimental protocol, the mice underwent this test thrice on a given day. Data for each mouse are expressed as the average values of three trials.

#### Forelimb grip assessment

A rat/mouse grip strength meter (SA415, Sansbio Co. Ltd., China) was used to assess the forelimb grip strength of the mice, which could be used to evaluate neuromuscular performance after ICH. Mice were held by their tails and allowed to grasp the steel grid with their forepaws. They were then pulled backward by their tails until they were pulled away from the grid. Standardized forelimb grip strength was expressed as forelimb strength measured in grams by the device divided by the mouse’s body weight in grams. The average values of three consecutive repeated assessments for each mouse were used for the analysis.

#### Hot-plate test

A hot-plate test was performed using a Hot/Cold Plate Video Analysis System (SA705, Sansbio Co. Ltd., China) to evaluate sensitivity to thermal stimuli. Mice were placed on a 52.0±0.3°C hot plate, and the latency of the first response (foot shake or paw lick) in the hind paw contralateral to the hematoma was recorded.

#### Balance beam test

A beam-walking apparatus (SA112, Sansbio Co. Ltd, China) was used to evaluate motor coordination and balance. The balance beam (125×2 cm) test was performed before surgery and at 7/14/21/28 days after ICH. All the mice were required to walk through the balance beam to reach a dark box (20×20×30 cm) located at the other end. The time required to enter the dark box was recorded. Each mouse was tested three times per day, and the average time (s) of the fastest 2 attempts was used for analysis.

### Brain tissue preparation

At 3/7/14/28 days after ICH, the mice were euthanized with 5% isoflurane and transcardially perfused with saline. For western blotting, brains were rapidly removed and cut into 1 mm-thick coronal sections using Mouse Brain Matrices. Target tissues surrounding the hematoma, including the caudate putamen (CPu), CC, globus pallidus, and internal capsule, were carefully dissected from sections and snap-frozen in liquid nitrogen. For luxol fast blue (LFB), transferase dUTP nick-end labeling (TUNEL), and immunofluorescence staining, mice were perfused followed with 4% paraformaldehyde (PFA) in 0.1 M phosphate-buffered saline (PBS) in 4°C. The brains were removed from the skulls, fixed in 4% paraformaldehyde for 24 hours, dehydrated in 30% sucrose solution for 48 hours, embedded in OCT Compound, snap-frozen for 1 h, and coronally sectioned in a series of 10 μm on a cryostat (CM1950, Leica). Sections were stored -20°C and in preparation for histological and immunofluorescence evaluations.

### Pathology

#### Luxol fast blue

To clarify the extent of myelin loss and histological changes in the white matter (WM) surrounding the hematoma after ICH, LFB staining was performed. The sections were stained with LFB (G1030, Servicebio) and eosin (G1002, Servicebio). Histological slides were scanned using a Pannoramic MIDI II scanner (3D Histech). We further evaluated WMI in the perihematomal CPu and CC after ICH using Case Viewer 2.4 (3D Histech, Hungary).

#### TUNEL

To estimate the number of apoptotic cells in the perihematomal region 3 and 7 days after ICH, a terminal deoxynucleotidyl TUNEL assay kit (G1501, FITC, Servicebio) was used according to the manufacturer’s instructions. Similarly, the apoptosis of OLs in the perihematomal region after ICH was evaluated by co-labeling with Olig2. The images were acquired using a panoramic MIDI II scanner. Four fields of view around the hematoma were selected in each section. TUNEL-positive cells were counted using ImageJ at 40× magnification; approximately 300–400 cells per mouse were analyzed.

#### Immunofluorescence

After blocking with 5% donkey serum for 1 h, slides were incubated with the primary antibodies overnight at 4°C, followed by washing thrice with PBS and appropriating for 1 h at 24℃ for 50 minutes in dark conditions. Nuclei were then incubated with DAPI (1:1000, G1012, Servicebio) solution at room temperature for 10 min in the dark. The following primary antibodies were used: rabbit anti-oligodendrocyte transcription factor 2 (Olig-2, 1:100, ab109186, Abcam), mouse anti-adenomatous polyposis coli (APC, 1:50, ab40778, Abcam), mouse anti-nerve/glial-antigen 2 (NG2, 1:100, ab259324, Abcam), mouse anti-Ki67 (Ki67, 1:50, ab15580, Abcam), mouse anti-myelin basic protein (MBP, 1:200, GB12226, Servicebio), mouse anti-ionized calcium binding adapter molecule 1 (Iba-1, 1:500, GB12105, Servicebio), rabbit anti-Fc RII/III receptor (CD16/CD32, 1:200, ab223220, Abcam). Secondary antibodies labeled with FITC and Cy3 were purchased from Service Bio. Immunofluorescent images were captured using a panoramic MIDI II scanner. In sections immunostained for Olig2, APC, NG2, Ki67, Iba-1, and CD16/32, the number of immunopositive cells (cells/mm^2^) was counted and analyzed using ImageJ software (version 1.54g) by an investigator blinded to the group and outcome assignments.

### Transmission electron microscopy

Ultrastructural observation of the WMI within the perihematomal CC after ICH was performed using transmission electron microscopy (TEM). Anesthetized mice were perfused intracardially with 0.1M PBS followed by 4% glutaraldehyde in 0.1 M PBS. Brain tissue 0 to 1 mm anterior to the bregma was cut into 1 mm-thick coronal sections using Mouse Brain Matrices. Approximately 1 mm^3^ of the right CC superior to the hematoma was extracted and stored in TEM fixative (G1102, Servicebio) at 4°C for 24 hours. The tissue blocks were washed using 0.1 M PB (pH 7.4) thrice for 15 min each. The blocks were then post-fixed with 1% OsO_4_ in 0.1 M PB (pH 7.4) for 2 h at room temperature, followed by rinsing in 0.1 M PB (pH 7.4) thrice for 15 min each. The blocks were dehydrated, embedded, polymerized, and sliced into 60∼80 nm thin sections on an ultra-microtome (EM UC7, Leica, Germany). Finally, the sections were placed onto 150-mesh cuprum grids with a Formvar film and stained with 2% uranium acetate and 2.6% lead citrate. Images were captured using a TEM (HT7800, HITACHI, Japan). Three images (4000× magnification) from each mouse and at least 20 random axons per image were used to measure the diameter of the axons, number of myelinated or unmyelinated axons per 100 μm^2^, and g-radio value. The g-radio value was calculated from the ratio of inner axonal diameter to overall diameter of the axon plus myelin, representing myelin thickness relative to axon diameter, where a higher g-ratio indicates a thinner myelin thickness.

### Enzyme-linked immunosorbent assay (ELISA)

Perihematomal tissues collected from each group were homogenized and centrifuged, and the supernatant was collected for analysis. The concentration of inflammatory cytokines at 3 and 7 days after ICH, including tumor necrosis factor-α (TNF-α), interleukin-1β (IL-1β), interleukin-6 (IL-6), and interferon gamma (IFN-γ) were measured using the specific ELISA reagent kits (TNF-α, ab208348, IL-1β, ab210895, IL-6, ab222503, Abcam; IFN-γ, 88-7314-88, Thermo Fisher Scientific) following the manufacturer’s instructions.

### Western blot analysis

Total protein in the perihematomal tissues was collected by homogenization and quantified using a BCA kit (Beyotime, China). To further investigate the protective mechanism of Sephin1 by prolonging the integration stress response, ATF4 (1:500, GB15001, Servicebio), CHOP (1:300, GB115720, Servicebio), T-eIf2α (1:3000, GB11544, Servicebio), and P-eIf2α (1:500, ab32157, Abcam) were used for western blot analysis. Chemiluminescent substrates were used to visualize the target proteins and performed a semi-quantitative analysis.

### Co-culture of primary microglia and oligodendrocytes

OLs were co-cultured with microglia activated by hemoglobin. Primary microglia and OLs were purchased from Warner Biotech (Wuhan, China). Primary microglia and OLs were incubated at 37°C and 5% CO_2_ for 24 h, and then gently seeded in a 24-well Transwell system (5×10^4^ cells/well and 1×10^5^ cells/well, respectively). Wells containing OLs were randomly assigned to two groups with or without Sephin1 (50 μM) in the culture medium. Hemoglobin (20 μM) was added to microglia while Sephin1 or vehicle was added to OLs for 6 h. The microglial-conditioned media were then transferred from the microglial cultures to the oligodendroglial cultures using a Transwell system. The CCK8 assay for the cell viability of OLs were measured after the cells interacted for 3/6/12/24 h. CCK-8 reagent was added into each well and incubated for an additional 2 h at 37°C. The optical density at 450 nm (OD450) was measured using a microplate reader, and each group was analyzed in triplicate.

### Statistical analyses

Statistical analyses were performed using SPSS 19.0. All data are presented as mean ± SD. Single comparisons were performed using unpaired t-tests, and multiple comparisons were performed using one-way or two-way analysis of variance (ANOVA). All tests were considered statistically significant at P<0.05.

## Results

### ICH leads to severe WMI

White matter and OL impairment is closely associated with poor neurological function after ICH.^11^ Based on the described method of inducing ICH, the scheme of the coronal brain section illustrates that the hematoma was localized below the CC and EC. The CPu and CC were set as the areas to evaluate WMI after ICH (Figure 1B). LFB staining revealed severe WMI in the CPu and CC surrounding the hematoma after ICH (Figure 1C and 1D). To quantify the changes in OLs in the Sephin1 and vehicle group after ICH, immunofluorescence staining for Olig2 and DAPI was performed. Three days after ICH, almost all OLs and other cells in the core region of the hematoma had completely disappeared. By day 7, OLs increased significantly and formed a ring around the hematoma in both ICH groups (Figure 1E).

### Sephin1 alleviated WMI and improved the long-term neurological function after ICH

MBP immunofluorescence staining revealed significant WMI in the perihematomal CPu and CC after ICH (Figure 2A). Damaged myelin was present in both ICH groups, and a myelin sheath with more severe damage was observed adjacent to the hematoma. Contrastingly, the ICH+Sephin1 group showed an obvious increase in MBP immunoreactivity in the CPu and CC groups compared to that in the vehicle group. Accordingly, Sephin1 mitigated WMI after ICH.

**Figure 2.**
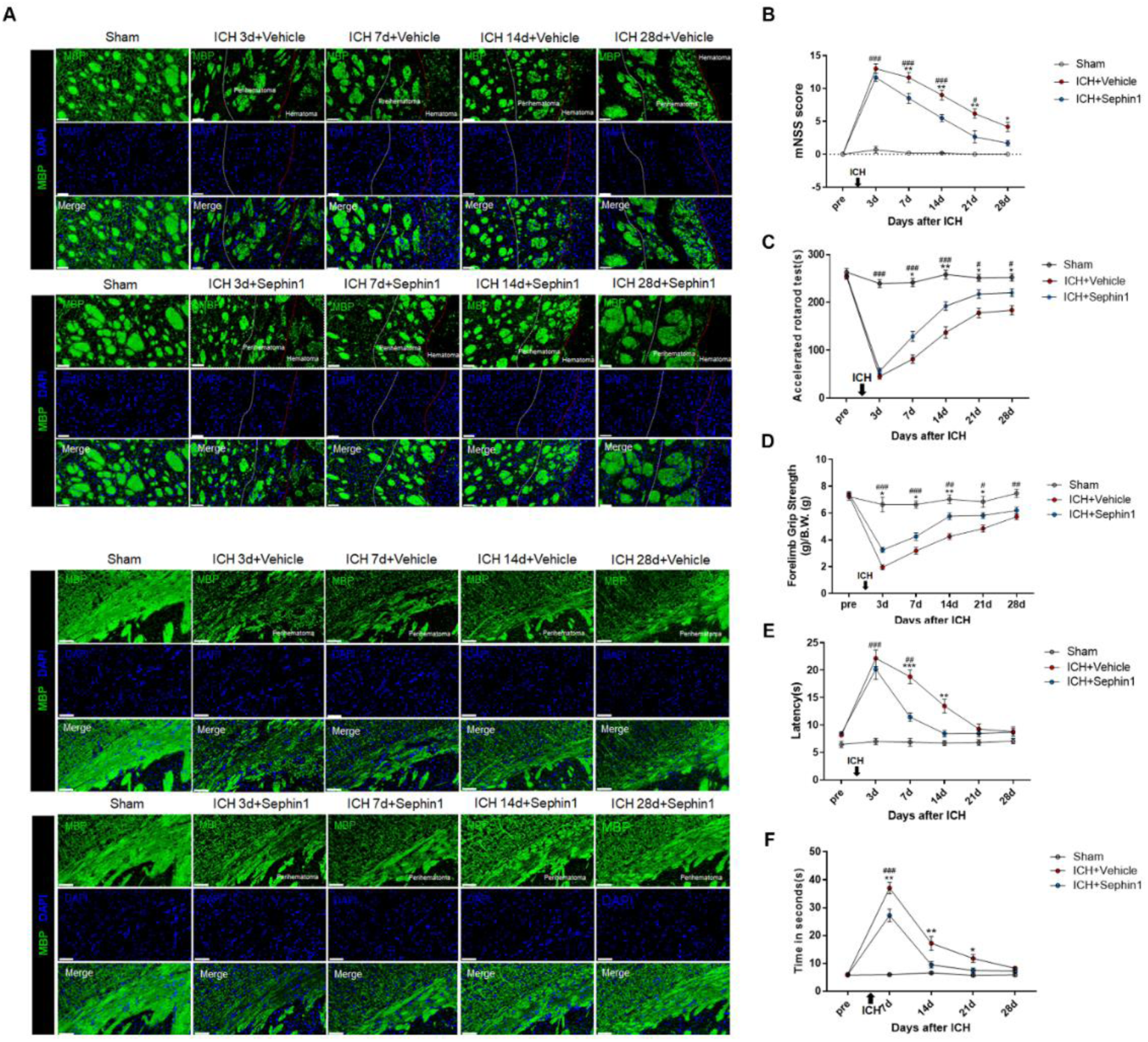
Sephin1 alleviated WMI and improved the long-term neurological function after ICH. **A,** Representative images of the MBP immunofluorescence staining in the perihematomal CPu (above) and CC (below). Lens: × 300; Scale bar: 50 μm. **B,** The neurological deficits of mice were assessed by mNSS. n=6 per group. **C,** The sensorimotor coordination and fatigue resistance of mice was assessed by accelerated rotarod test. n=5 per group. **D,** The forelimb grip strength of mice was measured by forelimb grip assessment. n=8 per group. **E,** The sensitivity of mice was evaluated by hot plate test. n=6 per group. **F,** The motor coordination and balance of mice was assessed by balance beam test. n=8 per group. ^#^*P* < 0.05, ^##^*P* < 0.01, ^###^*P* <0.001 versus Sham; \**P* < 0.05, \*\**P* < 0.01, \*\*\**P* < 0.001 versus ICH + Vehicle.

These outcomes of behavioral testing demonstrated that Sephin1 improved long-term neurological function after ICH. The highest mNSS score was observed 3 days after ICH, indicating that ICH contributed to severe neurological impairment. Consistent with the role of Sephin1 in mitigating ICH-induced WMI, the mNSS scores of mice in Sephin1 group were significantly lower than those in the vehicle group at 7/14/21/28 days after ICH (Figure 2B).

The rotarod test revealed that the sensorimotor coordination and fatigue resistance of Sephin1-treated mice was significantly improved compared to that of vehicle-treated mice (Figure 2C). Sephin1-treated mice showed significant and sustained enhancement of standardized forelimb grip strength (Figure 2D). Noticeable sensory deficits caused by ICH were alleviated in Sephin1-treated mice at 7 and 14 days after ICH (Figure 2E). In balance beam test, Sephin1-treated mice displayed a significant reduction in the time taken to enter the dark box compared to vehicle-treated mice at 7/14/21 days (Figure 2F).

### Sephin1 enhanced remyelination in the perihematomal region

TEM was performed to investigate the ultrastructural changes in axons and to measure the thickness of the myelin sheaths in the CC surrounding the hematoma. The axons in sham group were intact, with a compact myelin sheath and regular organelles. Conversely, swollen, demyelinated, or ruptured axons; non-compact or damaged myelin sheaths; and enlarged or vacuolated mitochondria were observed in ICH groups, and these symptoms significantly improved in Sephin1 group (Figure 3A). The number of myelinated axons and thickness of the myelin sheath dramatically decreased after ICH, which was remarkably improved by Sephin1 administration (Figure 3B-D). Quantification of the g-ratio was much lower in Sephin1-treated mice than in vehicle-treated mice 14 and 28 days after ICH (Figure 3E). Additionally, scatter plots of the axon diameters across the corresponding myelin thickness values revealed that Sephin1 greatly increased myelin thickness in the Sephin1 group compared to the vehicle group (Figure 3F).

**Figure 3.**
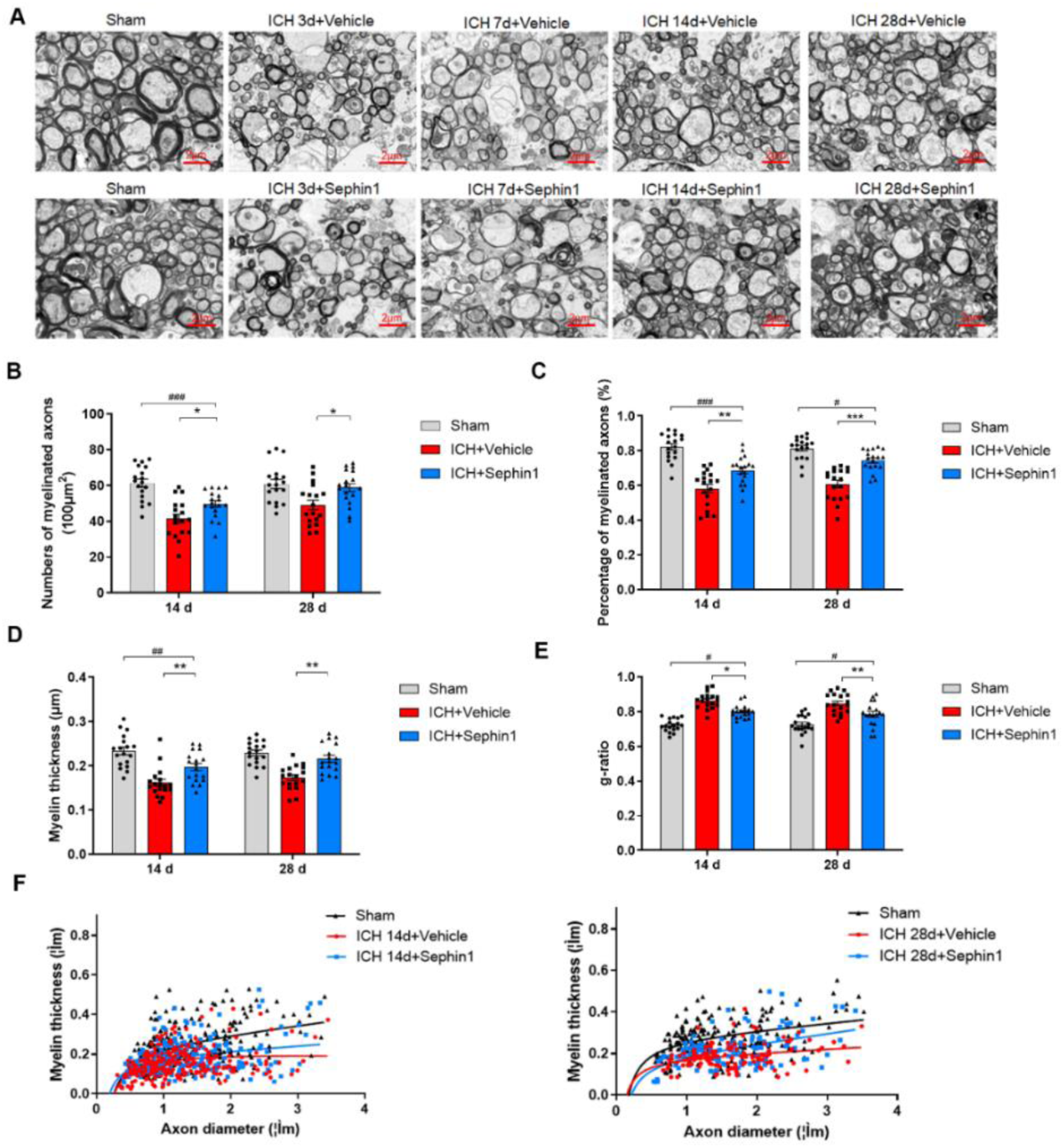
Sephin1 promoted remyelination in the CC surrounding the hematoma after ICH. **A,** Representative transmission electron microscopy images in the CC of mice from Sham, ICH + Vehicle and ICH + Sephin1 group after ICH. Lens: × 4000; Scale bar: 2 μm. **B,** Density of myelinated axons after ICH. n=18 per group. **C,** Quantification of the proportion of myelinated axons among total axons after ICH. n=18 per group. **D,** Quantification of the myelin thickness after ICH. n=18 per group. **E,** g-ratio of myelinated axons after ICH. n=18 per group. **F,** Scatter plot of myelin thickness values across the corresponding myelin thickness values at 14 and 28 days after ICH in Sham group (n=250 and 200 axons), ICH + Vehicle group (n=250 and 129 axons) and ICH+Sephin1 group (n=255 and 126 axons). ^#^*P* < 0.05, ^##^*P* < 0.01, ^###^*P* <0.001 versus Sham; \**P*< 0.05, \*\**P* < 0.01, \*\*\**P* < 0.001 versus ICH + Vehicle.

### Sephin1 increased the number of OLs in the perihematomal region

We quantified the temporal changes in the number of OLs in CPu and CC surrounding the hematoma from 3 to 28 days after ICH by immunofluorescence staining for Olig2 and DAPI. The density of Olig2^+^ cells in the peri-hematoma increased significantly in both ICH groups at 7 days after ICH, and yet progressively decreased at 14 and 28 days (Figure 4A and 4B). The number of OLs was significantly elevated in the CPu and CC of Sephin1-treated mice compared to the vehicle-treated mice at 14 and 28 days after ICH.

**Figure 4.**
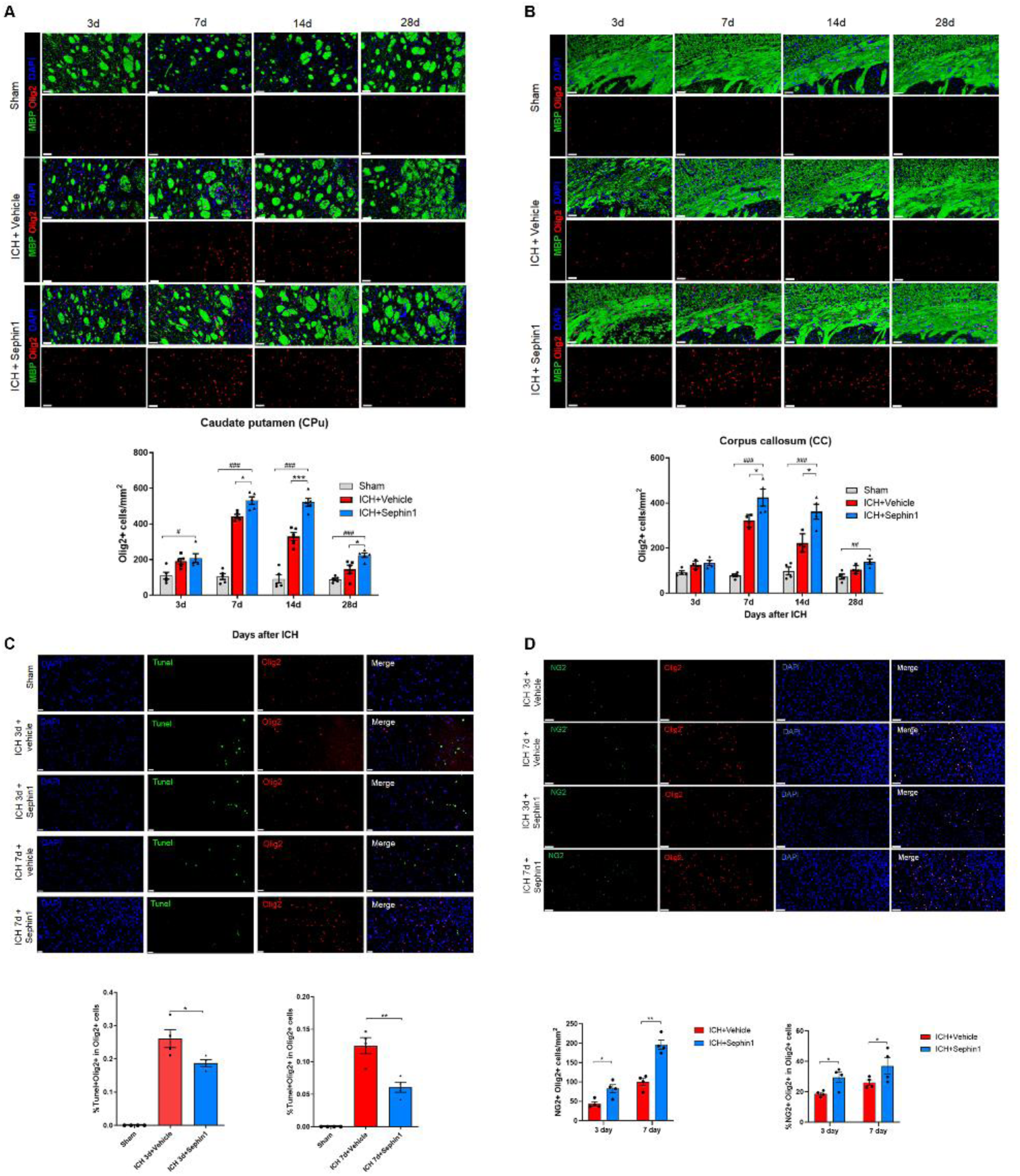
Sephin1 increased the number of OLs in the perihematomal region. **A,** Representative images of Olig2 and MBP immunofluorescence staining from the CPu after ICH. Lens: × 300; Scale bar: 50 μm. Quantitative analysis of Olig2+ cells in the region of CPu was performed. n=5 per group. **B,** Representative images of Olig2 and MBP immunofluorescence staining from the CC after ICH. Lens: × 300; Scale bar: 50 μm. Quantitative analysis of Olig2+ cells in the region of CC was performed. n=5 per group. **C,** Representative images of Tunel and Olig2 immunofluorescence staining in the perihematomal CPu. Lens: × 400; Scale bar: 20 μm. Quantitative analysis of the proportion of Tunel+Olig2+ cells among total OLs at 3 and 7 days was performed. n=6 per group. **D,** Representative images of NG2 and Olig2 immunofluorescence staining in the perihematomal CPu. Lens: × 300; Scale bar: 50 μm. Quantitative analysis of the number of OPCs and the proportion of OPCs among total OLs were performed. n = 4 per group. ^#^*P* < 0.05, ^##^*P* < 0.01, ^###^*P* <0.001 versus Sham; \**P* < 0.05, \*\**P* < 0.01, \*\*\**P* < 0.001 versus ICH + Vehicle.

Apoptosis, which is the primary pathway of OLs death, can be suppressed by prolonging ISR.^17^ After immunofluorescence staining, numerous TUNEL^+^Olig2^+^ cells were observed in the perihematomal CPu of all ICH mice (Figure 4C). ICH mice treated with Sephin1 exhibited a decrease in the proportion of TUNEL+Olig2+ cells to Olig2^+^ cells at 3 and 7 days after ICH compared to vehicle-treated mice, indicating that Sephin1 provides potent protection for OLs. There was no significant difference in the population of TUNEL^+^Olig2^+^ cells between vehicle- and Sephin1-treated mice at 3 and 7 days after ICH (Supplementary Figure 2A).

OPC have the capacity to proliferate and differentiate into mature OLs.^12^ In the perihematomal region, NG2 and Olig2 co-labeled cells were used to identify OPCs. The population and proportion of OPCs in OLs was significantly increased in Sephin1-treated mice versus vehicle-treated mice (Figure 4D). Additionally, Ki67 and Olig2 double immunofluorescence staining was used to identified proliferating OLs. We investigated whether the increased number of proliferating OLs around the hematoma primarily migrated from the subventricular zone (SVZ), which is the site of oligodendrogenesis. We found that the number of proliferating OLs in the ipsilateral SVZ was more than 10-fold lower than that in the peri-hematomal area on days 3 and 7 after ICH (Supplementary Figure 2B). The proportion and population of proliferating OLs in Sephin1-treated mice were significantly higher than those in the vehicle group at 3 and 14 days post-ICH (Supplementary Figure 2C). We also found a slight increase in the number of mature OLs (APC^+^Olig2^+^ cells) around the hematoma in mice treated with Sephin1 compared to that in mice treated with vehicle; however, there was no significant difference in the proportion of mature OLs among the total OLs (Supplementary Figure 2D).

### Sephin1 modulated microglial polarization and decreased the release of pro-inflammatory cytokines

After ICH, pro-inflammatory molecules released from neurons and other cells result in microglia activation.^30^ Activated microglia are polarized into the classic M1 (pro-inflammatory) or alternative M2 (anti-inflammatory) phenotypes, with M1 microglia exacerbating OL death and WMI after ICH.^31^ Compared to the vehicle group, the number of CD16/CD32^+^ Iba1^+^ cells significantly decreased in Sephin1-treated mice at 3 and 7 days after ICH, suggesting that the administration of Sephin1 modulated the polarization of microglia/macrophages (Figure 5A and 5B). At 3 and 7 days after ICH, the expression levels of IL-6, IL-1β, TNF-α, and IFN-γ in perihematomal tissues were significantly increased in both ICH groups compared with the sham group. Compared with the mice in vehicle group, the expression levels of IL-6, IL-1β, TNF-α, and IFN-γ were significantly decreased in Sephin1 group (Figure 5C-F).

**Figure 5.**
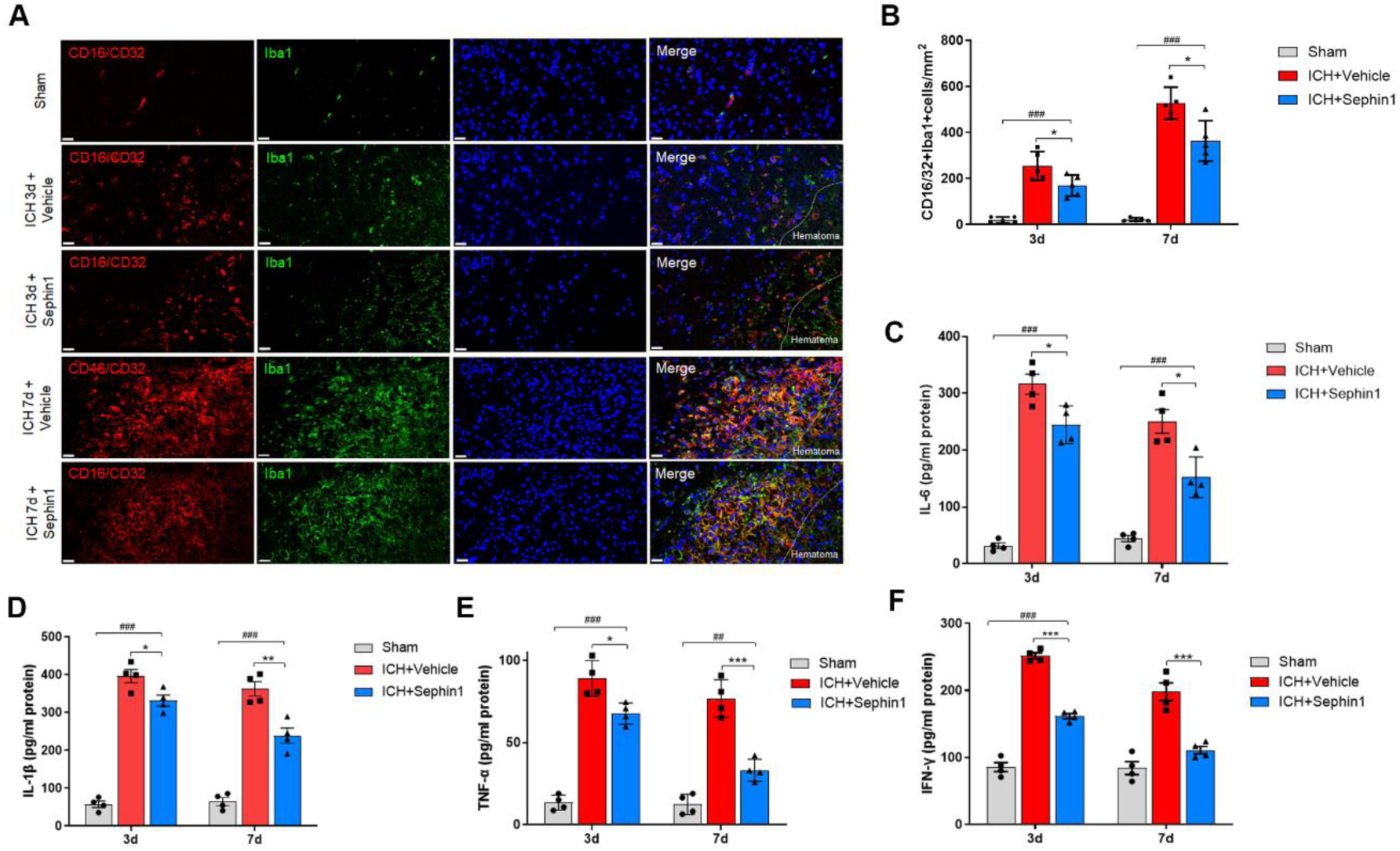
Sephin1 modulated the polarization of microglia and reduced the neuroinflammation after ICH. **A,** Representative images of CD16/32 and Iba1 double immunofluorescence staining at 3 and 7 days post ICH. Lens: × 400; Scale bar: 20 μm. **B,** Quantification of CD16/32+Iba1+ cells in perihematomal region. n=5 per group. **C,** Quantitative analysis of expression level of IL-6 was performed. n=4 per group. **D,** Quantitative analysis of expression level of IL-1β was performed. n=4 per group. **E,** Quantitative analysis of expression level of TNF-α was performed. n=4 per group. **F,** Quantitative analysis of expression level of INF-γ was performed. n=4 per group. ^##^*P* < 0.01, ^###^*P* <0.001 versus Sham; \**P* < 0.05, \*\**P* < 0.01, \*\*\**P* < 0.001 versus ICH + Vehicle.

### Molecular mechanism of Sephin1 on protecting OLs

The activation of the PERK/eIF2α/CHOP pathway of ISR acts as a regulator for cell apoptosis.^19^ Here, the activation of ISR-related signaling pathways was confirmed by western blot analysis of brain tissue lysates from the perihematomal region (Figure 6A). The expression levels of P-eIF2α were significantly different among the groups, with the highest levels observed in the ICH+Sephin1 group (Figure 6B). There were no significant changes in the levels of T-eIF2α among all groups. Moreover, quantitative analysis demonstrated significant inhibition of CHOP and ATF4 expression by Sephin1 treatment compared to vehicle-treated mice.

**Figure 6.**
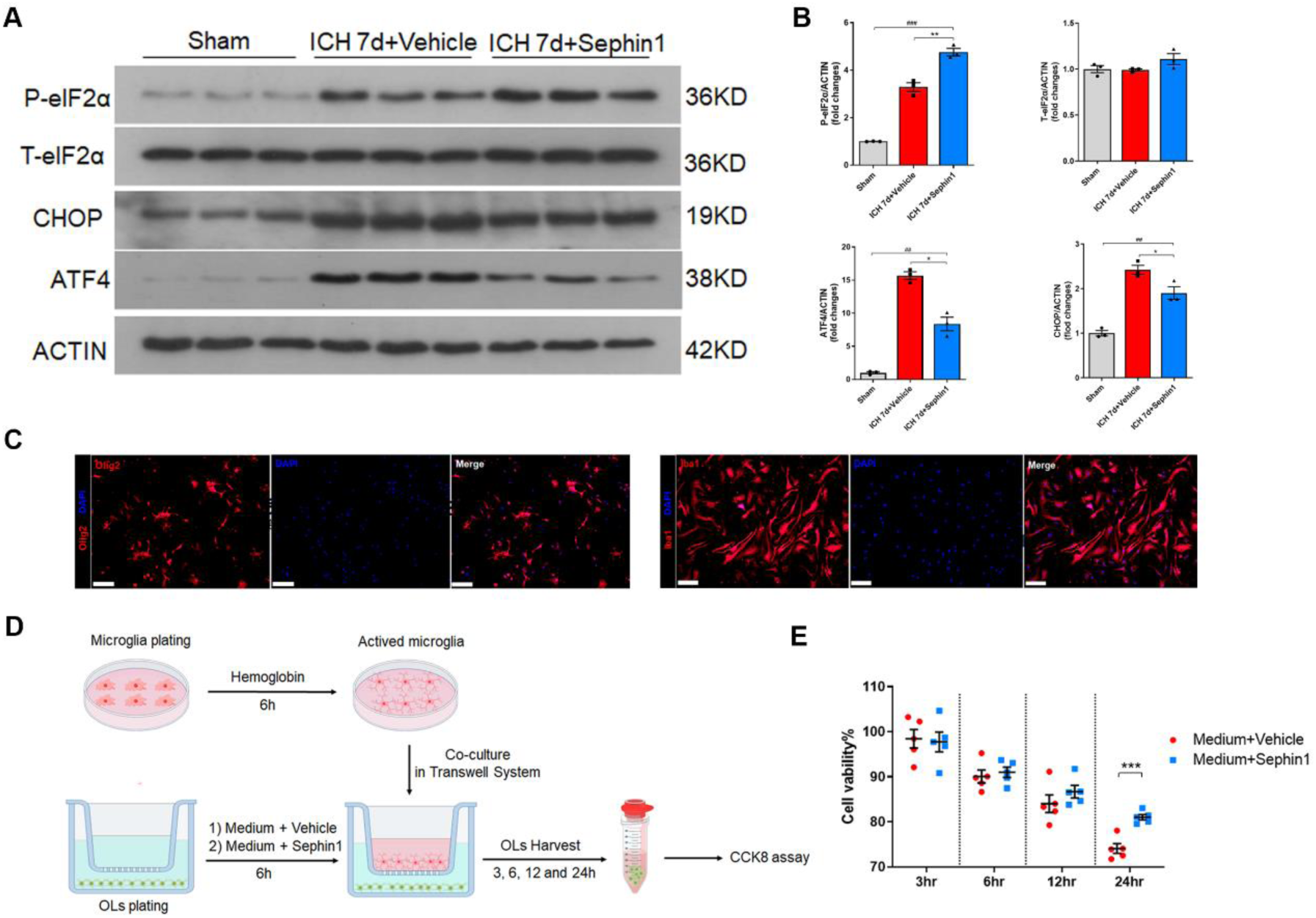
Sephin1 protected OLs when co-cultured with the activated microglia. **A,** Representative western blot bands showed the expression of the key components of the ISR, CHOP, ATF4, P-eIF2α and T-eIF2α in the perihematomal tissues at 7 days. **B,** Quantitative analyses of expression levels of P-eIF2α, T-eIF2α, CHOP and ATF in the perihematomal tissue at 7 days were performed. ^##^*P* < 0.01, ^###^*P* < 0.001 versus Sham; \**P* < 0.05, \*\**P* < 0.01 versus ICH + Vehicle. n=3 per group. **C,** Primary OLs was verified by staining with the oligodendroglial marker Olig2. Primary microglia was verified by staining with the microglial marker Iba1. Lens: × 200; Scale bar: 50 μm. **D,** Overview for the vitro portion of the present study. **E,** Cell viability of primary OLs after co-cultured with activated microglia was determined using CCK8 assays. \*\*\**P* < 0.001 versus Medium + Vehicle. n=5 per group.

Sephin1 prolongs the benefit of eIF2α phosphorylation in OPCs stimulated by IFN-γ *in vitro*.^27^ In present study, primary OLs and microglia stimulated by hemoglobin were co-cultured in a transwell system for 24h (Figure 6C and 6D). The cell viability of OLs treated with Sephin1 was significantly higher than that of OLs treated with vehicle, implying that Sephin1 could protect OLs in pro-inflammatory environments (Figure 6E).

## Discussion

In the present study, we identified the effects of Sephin1 in protecting OLs and ameliorating WMI in a mouse model of ICH-induced by injecting bacterial collagenase VII-S. Sephin1 significantly alleviated WMI and promoted the remyelination of injured axons in the perihematomal region, paralleled by an improvement in long-term neurological function. Sephin1 also inhibits the apoptosis of OLs, promoted their proliferation of OLs, and increased the number of OLs in the perihematomal region after ICH. Sephin1 treatment inhibits microglial polarization to the M1 phenotype and decreased the release of pro-inflammatory cytokines in perihematomal tissues. In addition, Sephin1 treatment inhibits the expression of CHOP and ATF4 at 7 days after ICH and prolonged ISR. Taken together, these results support the idea that Sephin1 treatment may be beneficial in patients with ICH. WMI is closely associated with long-term neurological impairments and fatality in patients with ICH.^2,10,32^ Our results demonstrated that the white matter in the perihematomal region was damaged after ICH, manifested as swollen, demyelinated, or ruptured axons, non-compact or damaged myelin sheaths, and enlarged or vacuolated mitochondria. ICH-induced WMI is attributed to a diverse range of pathological processes such as the direct compressive effect of the hematoma, neuroinflammation, coagulation cascade, metabolic products of the hematoma, and oxidative stress.^33,34^ However, effective therapeutic strategies to restore WMI caused by ICH are lacking. Therapies aimed at protecting OLs can promote recovery from WMI and improve neurological function after ICH.^35,36^ Our results suggested that Sephin1 attenuated WMI and significantly improved the neurological function of mice with ICH in a series of behavioral tests, consistent with previous studies showing that as histological damage to the white matter is mitigated, the corresponding neurological dysfunction is improved.^37,38^

Sephin1 shields OLs against inflammatory stress and improves neuropathy in various neurological diseases by prolonging the ISR.^19,24,39^ To date, there has been little understanding of the protective effects of Sephin1 on OLs in ICH. Therefore, we examined the apoptosis and differentiation of OLs, which may impact the number and function of OLs.

OLs, especially OPCs that are susceptible to apoptosis and necrosis after ICH, are critical for remyelination during WMI.^40^ Sephin1 treatment increased the number of OLs in the perihematomal region 7 and 14 days after ICH, which might promote remyelination and alleviate WMI. A substantial proportion of apoptotic OLs at 3 and 7 days after ICH, which was significantly decreased in the ICH+Sephin1 group. Furthermore, we observed that Sephin1 treatment increased the proliferative capacity of OLs around the hematoma but did not alter the proportion of mature OLs, which implied that Sephin1 might not dramatically affect the maturation of OLs. However, the exact mechanism by which Sephin1 affects OL proliferation and maturation remains unclear. This is likely owing to the fact that Sephin1 protects OLs mainly by prolonging the ISR and inhibiting global protein synthesis. Additionally, co-culture of OLs with activated microglia confirmed the protective effect on cell viability of Sephin1 by prolonging ISR. Therefore, Sephin1 protects OLs against pro-inflammatory environments in a manner independent of immunomodulation and crosstalk with other cells.

Secondary neuroinflammation in the perihematomal region after ICH, including the activation of M1 microglia and the secretion of pro-inflammatory cytokines, is an important factor in the exacerbation of WMI, which is also considered to hinder both remyelination and white matter repair.^41,42^ We examined M1 polarization of microglia and measured pro-inflammatory cytokines, including IL-6, IL-1β, TNF-α, and IFN-γ, in the perihematomal tissues by ELISA. The data showed that Sephin1 treatment reduced the number of M1 microglia and inhibited pro-inflammatory cytokine secretion 3 and 7 days after ICH.

Inhibiting the M1 polarization of microglia could attenuate WMI and improve neurological function after ICH.^33,43^ However, the specific mechanism by which Sephin1 modulates microglial recruitment and polarization requires further investigation.

Excessive activation of ISR in stressed cells leads to the enhanced translation of the transcription factors ATF4 and CHOP, causing up-regulation of GADD34 expression, which in turn results in the dephosphorylation of eIF2α and restores protein synthesis.^17^ An increased rate of protein synthesis promotes the accumulation of protein aggregates, ultimately inducing apoptosis of stressed cells.^44^ Notably, the pharmacology of Sephin1 has been confirmed to have no effect on non-stressed cells, with short half-lives in organs and plasma of less than 6 h.^26,45^ Moreover, GADD34 mutant mice, which lack GADD34-PP1R15A activity, showed normal myelination in the presence of persistent neuroinflammation.^46^ Taken together, these data suggest that cells in perihematomal tissues, such as OLs, are likely tolerant to the transient pharmacological inhibition of GADD34 activity caused by Sephin1.

Our study has several limitations. In the present study, we identified the effect of Sephin1 on the inhibition of OLs apoptosis in mice with relatively mild ICH, whereas the effect in a severe ICH model is not clear yet. We could not confirm whether the current dose of Sephin1 would exert the equivalent effect in a severe ICH model. In addition, we did not explore the effects of Sephin1 treatment on neurons and astrocytes, which might play crucial roles in the pathophysiology of ICH. It is well known that neurons are closely associated with neurological function after ICH; similarly, astrocytes are highly involved in neuroinflammation and neurotoxicity.^47,48^ Therefore, further investigation of the optimize dosing of Sephin1 and the effects on diverse cell types in more clinically relevant models of ICH is needed in the future.

## Conclusion

We demonstrated that treatment with Sephin1 effectively protects OLs and ameliorates WMI, contributing to improved long-term neurological function in a mouse model of ICH. The mechanisms of protection of OLs with Sephin1 may involve inhibition of protein synthesis, modulation of neuroinflammatory responses, and reduction of apoptosis after ICH. Given the significant effect of WMI on ICH prognosis, our findings suggest that Sephin1 treatment is a promising therapeutic strategy for ICH.

## Acknowledgements

Not applicable.

## Supplemental Material

Figure S1-S2

